# Wnt and Fgf signaling pharmacological inhibition affect posterior growth during *Tribolium castaneum* germband elongation

**DOI:** 10.1101/2025.03.14.643200

**Authors:** Marco Mundaca-Escobar, Renato V. Pardo, Rodrigo E. Cepeda, Andres F. Sarrazin

## Abstract

Axial elongation and sequential segmentation are developmental processes that occur simultaneously and are highly conserved in vertebrates and most arthropods. These features rely on the dynamic expression of a genetic network that establishes the segmented patterning and regulates various cellular behaviors, including tissue rearrangements and cell divisions. In vertebrates, Wnt and Fgf signaling are essential for these processes. While some studies in arthropods have linked these pathways to segmentation, there is still much to discuss regarding their regulatory role in cellular processes. In this study, we pharmacologically inhibited Wnt and Fgf signaling pathways by exposing developing *Tribolium castaneum* embryos to IWP-3 and SU5402, respectively. We observed that both treatments resulted in a shortening of the embryos and a decrease in the number of cell divisions during a period characterized by high proliferation rates. Although the segmented patterning was not disrupted, the segments were smaller in the embryos treated with the Fgf inhibitor than in the controls. Additionally, time-lapse imaging revealed that cell movement along the anteroposterior axis was affected in the IWP-3-treated embryos. In contrast, Fgf inhibition primarily altered the direction of cell movements at the posterior end of the embryo. Our findings provide insight into the roles of Wnt and Fgf signaling pathways in regulating significant cellular behaviors during the posterior growth of *Tribolium* and possibly other arthropods.

## Introduction

A segmented body plan is a shared characteristic of chordates and arthropods. During development, all vertebrates and most arthropods add their segments sequentially while the main body axis elongates, in a phenomenon known as posterior growth [1, 2]. This process relies on a relatively conserved gene regulatory network to coordinate the establishment of the segmented patterning and the different cellular behaviors that allow the body to extend along its anteroposterior axis [3–5]. In vertebrates, it has been defined that Notch, Wnt, and Fgf signaling are essential for axis elongation and posterior segmentation, even though there are significant differences between the different organisms (*i.e.*, zebrafish, chicken, and mouse) analyzed [6–14]. Some studies have also linked these signaling pathways (although Fgf to a lesser extent) [see 15, 16] to posterior growth in sequentially segmented arthropods [reviewed in 3, 5, 17, 18]. However, little has been delved into their specific regulatory roles during segmentation, elongation, and the cellular processes involved.

Like vertebrate somitogenesis, sequential segmentation in arthropods is governed by a segmentation clock [19–22], where a molecular oscillator generates a temporal periodicity that is progressively translated to a spatially repeated pattern. Until now, the oscillating genes involved in arthropod body segmentation have been discovered within the “pair-rule” genes family [19, 20, 22–27] or the Notch/Delta signaling pathway [19, 24, 26–28], depending on the studied species. Along the embryo, the very dynamic expression of those cyclic genes is restricted to a posterior region called the segment addition zone (SAZ) that is preceded anteriorly by the stable expression of the segment polarity genes (*e.g.*, *wingless*, *engrailed*, *hedgehog*) at the site (the wavefront) where oscillations stop, and new segments arise. In vertebrates, posterior Wnt/Fgf signaling gradients mainly define this wavefront, keeping more posterior cells undifferentiated while oscillating [29]. It has been proposed in arthropods that the Wnt signaling pathway organizes posterior growth and maintains the genetic oscillations through the regulation of *caudal*, a highly conserved transcription factor expressed as a posterior gradient in the SAZ [reviewed in 5, 18]. In various arthropods, the knock-down of different components of Wnt signaling severely affects segmentation. For example, in the cricket and the milkweed bug, blocking *armadillo*/β-catenin or *pangolin*/TCF expression generates truncated embryos without posterior segments [30, 31]. In *Tribolium*, Wnt1/*wg* loss-of-function embryos are shorter and have smaller segments, as well as RNAi against Wnt8/D, both in the spider *Parasteatoda* and *Tribolium*, generates malformed abdominal segments and a smaller SAZ [32, 33]. Similar results were obtained by knocking down Wnt receptors Frizzled1 and Frizzled2 in *Tribolium*, impairing axial elongation and partially eliminating the SAZ [16]. As well as posterior gene oscillations, the maintenance of undifferentiated cells with the capacity to proliferate at the posterior part of the embryo is most probably also regulated by the Wnt-dependent gradient of *caudal* expression [34, 35]. Reduced cell proliferation within the SAZ of the cockroach *Periplaneta* was found after using RNAi against Wnt1 and by pharmacological treatment with the inhibitor of the Wnt signaling pathway, IWP-3 [36]. In addition, using RNA-seq after RNAi against Wnt/β-catenin pathway members, Oberhofer et al. (2014) [35] identified several Wnt downstream genes required in the SAZ for patterning, mesoderm formation, and cell division.

It is clear from the above-mentioned Wnt functional analyses that this signaling pathway must be linked to cellular processes as convergent extension movements and/or cell proliferation that are already well-known driving forces during posterior growth in sequentially segmenting arthropods [21, 37–42]. Therefore, the principal aim of this study was to unravel the regulatory role that the Wnt signaling pathway has on the cellular processes associated with posterior growth, in addition to the potential participation of the Fgf signaling pathway, which has been less studied in arthropod segmentation but is known to be crucial in vertebrate somitogenesis along with the Wnt pathway. Using dissected *Tribolium* embryos for pharmacological manipulation, we found shorter embryos with fewer cell divisions after inhibiting both signaling pathways. Although the segmentation process itself was not affected, Fgf signaling inhibition showed smaller segments compared to control. In addition, fluorescent *in vivo* imaging showed that cell motion along the anteroposterior axis was affected when we blocked Wnt signaling. At the same time, Fgf inhibition mainly altered the direction of cell movements at the posterior rear of the embryo. Our findings suggest that Wnt and Fgf signaling pathways are essential for regulating cell proliferation and cellular movements, both cellular behaviors with a prominent role during *Tribolium* and other arthropods’ posterior growth.

## Materials and methods

Insect husbandry, whole embryo culture, pharmacological manipulation, single-embryo *in situ* hybridization, and embryo length measurement were carried out as described in Macaya et al. (2016) [43]. Phospho-Histone H3 pSer10 antibody staining and the subsequent proliferating cell quantification were performed essentially as in Cepeda et al. (2021) [40]. For the pharmacological inhibition of Wnt and Fgf signaling pathways, we tested different concentrations of inhibitors: IWP-3 (Inhibitor of Wnt production-3; Sigma SML0533) at 0.45 µM, 4.5 µM, 22.5 µM and 45µM, and SU5402 (Fgf receptor inhibitor; Sigma SML0443) at 0.5 µM, 2.5 µM, 5 µM and 50 µM. Based on our statistical analysis of the effect on embryo length (S1 Fig), we selected the lowest effective concentrations for subsequent analyses (45 µM working solution for IWP-3 and 2.5 µM for SU5402).

Additionally, we synthesized a *Tc-wnt1* digoxigenin-labeled RNA probe from a PCR amplified sequence (forward primer 5’-GGGATGTTACCAACGCCAGA-3’ and reverse primer 5’-CTTCACTTCACAGCAATG-3’) cloned into DH5α competent cells (ThermoFisher). For segment area analysis using *in situ* hybridization images, we measured the length and width at the center of each specific segment (Maxillary, Labial, Thoracic 1, and Thoracic 2), delimited by the rostral border expression of *Tc-wnt1*, using the ImageJ Straight Line tool.

### Embryo collection

All embryos used in this work were derived from the *Tribolium castaneum* EFA-nGFP transgenic line, which expresses nuclear-localized GFP throughout the body [21]. Embryonic stages were measured in minutes post horseshoe stage (mph), as defined in Cepeda et al. (2017) [39]. The horseshoe stage is characterized by the posterior amniotic fold extending anteriorly, forming a recognizable horseshoe-shaped amnion that overlies part of the germband [44]. All embryos were dissected at the 0-mph stage, following the procedures outlined in Macaya et al. (2016) [43], and were subsequently fixed in 4% formaldehyde at the desired stage, as required for each specific experiment.

### Cell movements analysis

Live imaging was conducted at 30°C using a Nikon Eclipse Ci-L microscope. All image analyses were performed using the free software ImageJ (FIJI). We utilized the ImageJ Extended Depth of Field plugin to generate a composite image of the Z-projection photographs. Each photograph was adjusted for brightness and contrast, and a movie was compiled. Using the Manual Tracking plugin, we tracked the individual fluorescent nuclei of dissected 0 mph EFA-nGFP embryos for 2 hours at 5-minute intervals. Cell movements (12 cells per embryo) from three different regions of the germband (*i.e.*, anterior, middle, and posterior) were mapped (Fig 4A). We measured three aspects of cell movement: i) total cell displacement, which is the sum of the distances covered by each cell over 2 hours, ii) net cell movement or effective cell displacement, which is the distance between the starting and ending points, and iii) the angle of the cell’s movement, which is the angle between the anteroposterior axis of the germband and the resulting displacement vector.

### Statistical analysis

The data were analyzed using GRAPHPAD PRISM v9.5.0 software. We assessed data distribution with the D’Agostino-Pearson test. To determine the statistical significance of germband length, cell proliferation, and segment area, we compared groups with a normal distribution using an unpaired *t*-test. For groups without a normal distribution, we employed the nonparametric Mann-Whitney test. For the analysis of cell movements in the films, a one-way anova TEST was performed to compare the variables and then a Tukey’s multiple comparisons test was performed. Asterisks indicate statistical significance: * *p* < 0.05, ** *p* < 0.01, *** *p* < 0.001, and **** *p* < 0.0001.

## Results

### Wnt and Fgf signaling pharmacological inhibition affect germband elongation and cell proliferation

To investigate the role of Wnt and Fgf pathways specifically on *Tribolium* germband elongation, we first exposed dissected fluorescently labeled EFA-nGFP embryos to the Inhibitor of <u>W</u>nt <u>p</u>roduction (IWP-3) that blocks Wnt palmitoylation by inactivating the O-acyltransferase Porcupine [36, 45], and the Fgf receptor inhibitor SU5402 [46], at different concentrations (S1 Fig). Next, we evaluated the effect of both chemicals on germband length and cell proliferation (PH3 positive cells normalized by germband area; see Experimental Procedures) during elongation, 2-hour from the onset of posterior growth (at the horseshoe stage) [see 43, 44]. After pharmacologically inhibiting both signaling pathways, statistically significant differences were obtained when we measured the length of the treated embryo at 45 µM for IWP-3 (*p* = 0.0353) and from 2.5 µM to 50 µM for SU5402 (*p* = 0.0061 to *p* < 0.0001, respectively), compared to controls (S1A and S1B Figs). A significant decrease in cell divisions was found at the same concentrations affecting embryo lengthening for both signaling inhibitors; 45 µM for IWP-3 (*p* = 0.001) and from 2.5 µM to 50 µM for SU5402 (*p* = 0.0011 to *p* < 0.0001, respectively) (S1C and S1D Figs). After the drug testing, we selected the lowest effective concentration for subsequent analyses, *i.e.*, 45 µM in the case of IWP-3 and 2.5 µM for SU5402. Thus, using these concentrations, we treated embryos with each inhibitor – and a combination of both – during increasing time intervals (1, 2, 3, and 4 hours after the onset of posterior elongation) and evaluated germband elongation (Fig 1 and S2 Fig) and cell proliferation (Fig 2 and S2 Fig).

**Fig 1.**
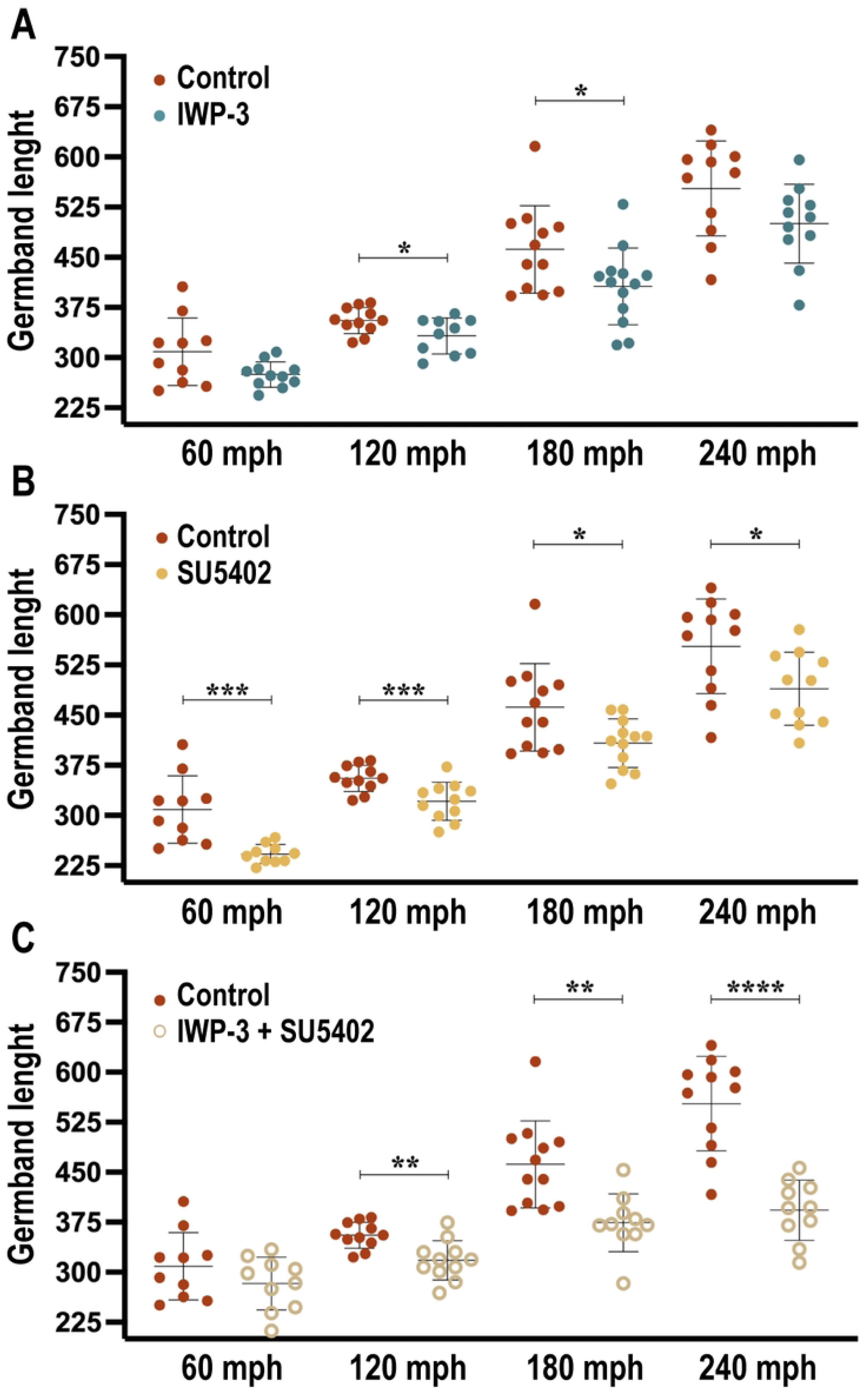
Germband shortening after Wnt and Fgf signaling pharmacological inhibition. (A-C) Dissected embryos were exposed to increasing time intervals, ranging from 60 to 240 minutes post horseshoe stage (mph; the onset of elongation), with 45 µM IWP-3 (A), 2.5 µM SU5402 (B), and a combination of both drugs (C). Each treatment was compared to an equivalent DMSO control incubation. The error bars indicate the standard deviation (SD) of the mean (*n* = 10 – 13). Asterisks indicate statistically significant differences according to the Mann-Whitney test and the Unpaired *t*-test. For IWP-3 treatment, * *p* = 0.0353 at 2 hours and *p* = 0.0335 at 3 hours. For SU5402 treatment, *** *p* = 0.0008 at 60 mph, ** *p* = 0.0035 at 120 mph, * *p* = 0.0212 at 180 mph, and *p* = 0.029 at 240 mph. For the combined IWP-3 + SU5402 treatment, ** *p* = 0.0021 at 120 mph, *p* = 0.0017 at 180 mph, and **** *p* = < 0.0001 at 240 mph.

**Fig 2.**
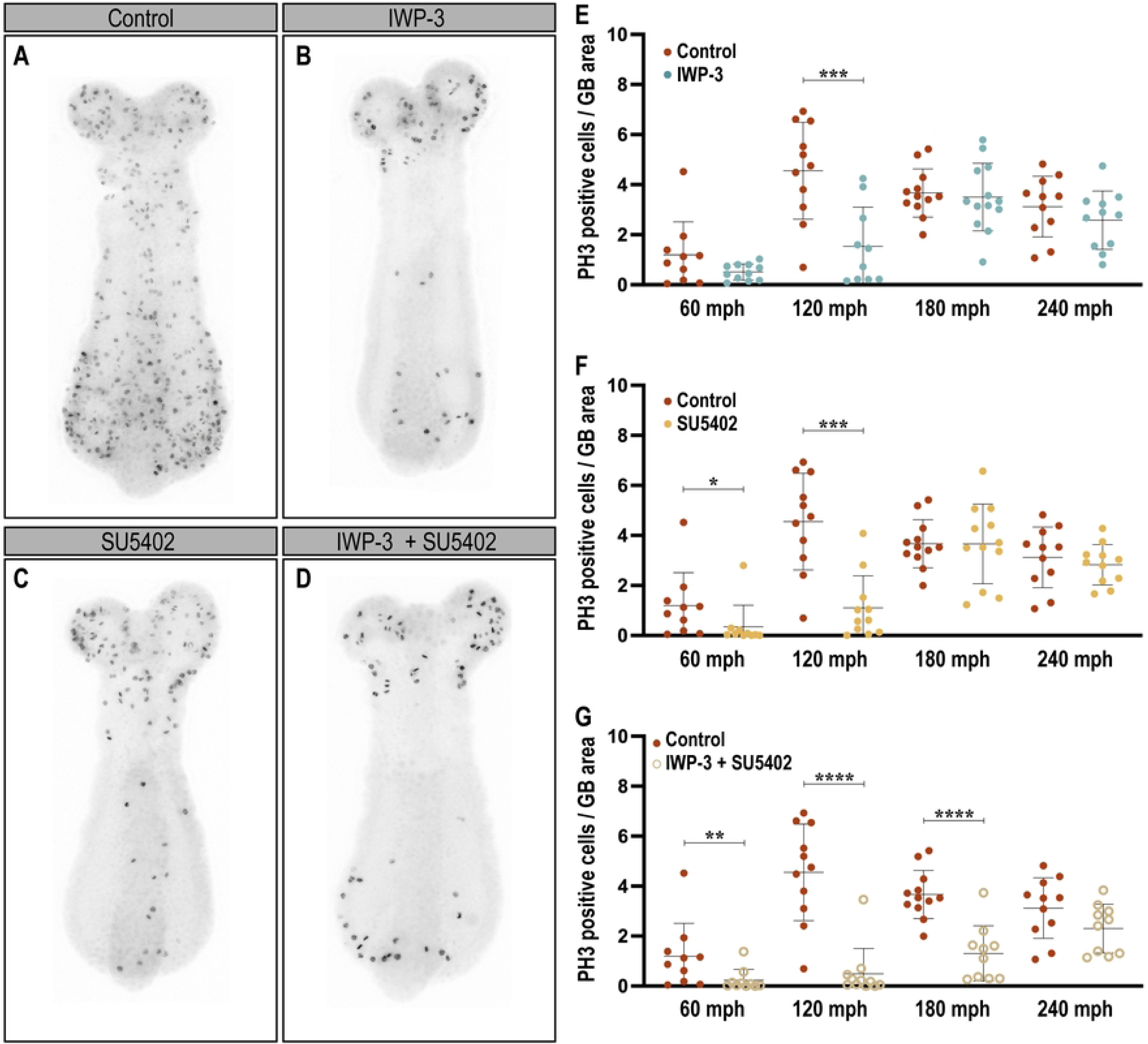
Reduced cell divisions after IWP-3, SU5402, and IWP-3+SU5402 incubations. (A-G) Dissected embryos were incubated during increasing time intervals, from 60 to 240 mph with DMSO (A, E-G), 45 µM IWP-3 (B, E), 2.5 µM SU5402 (C, F), and a combination of both drugs (D, G). Each treatment was compared to the equivalent DMSO control incubation. (A-D) Representative pictures of 120 mph PH3 stained dissected embryos incubated for 2 hours with DMSO (A; control), 45 µM IWP-3 (B), 2.5 µM SU5402 (C), and a combination of both chemicals (D). Red fluorescent antibody-stained embryo pictures were converted to black and white images and inverted to improve visualization. All embryos are shown in dorsal views. The anterior is to the top. The error bars indicate the SD of the mean (*n* = 10 – 13). Asterisks indicate statistically significant differences according to the Mann-Whitney test and the Unpaired *t*-test. For IWP-3 treatment, *** *p* = 0.001 at 120 mph. For SU5402 treatment, * *p* = 0,0146 at 60 mph, *** *p* = 0.0003 at 120 mph. For the combined IWP-3 + SU5402 treatment, ** *p* = 0.0088 at 60 mph, **** *p* = < 0.0001 at 120 mph, and **** *p* = < 0.0001 at 180 mph.

Using IWP-3, we found a significant shortening in the embryo length after 2- and 3-hour incubation (*p* = 0.0353 and *p* = 0.0335, respectively) but not after 4 hours (*p* = 0.0728) (Fig 1A). On the other hand, SU5402 incubation showed an effect on elongation after all time intervals evaluated (*p* = 0.0008, *p* = 0.0035, *p* = 0.0212 and *p* = 0.029 for 1-, 2-, 3-, and 4-hour incubations, respectively) (Fig 1B). Combining both inhibitors resulted in a length reduction from 2- to 4-hours incubation (*p* = 0.0021, *p* = 0.0017 and *p* <0.0001, respectively) (Fig 1C).

Strikingly, when we quantified the amount of PH3^+^ cells at each time point (Fig 2), we observed significant changes in cell proliferation only after 1- and 2-hour inhibiting Fgf signaling (*p* = 0.0146 and *p* = 0.0003, respectively) and 2-hour incubation with IWP-3 (*p* = 0,001) (Fig 2A-C, E, F). In addition, the combined treatment of 45 µM IWP-3 and 2.5 µM SU5402 during 1, 2, and 3 hours resulted in a statistically significant reduction of dividing cells along the germband (*p* = 0.0088, *p* < 0.0001 and *p* < 0.0001, respectively) (Fig 2D, G).

### Fgf signaling inhibition impact on the size of germband segments

To analyze proper segment formation on IWP-3 and SU5402 treated embryos, we performed *in situ* hybridization against *Tc-wnt1* mRNA. *Tc-wnt1* is expressed in the rostral border of each segment formed and in a patch of cells at the posterior tip of the embryo [32]. After using each inhibitor, the segmented patterning along the trunk and posterior region of the germband was not disrupted, but the striped expression of *Tc-wnt1* lost its definition (Fig 3A-C). Notably, the posterior expression remained unaltered compared to the control. Moreover, when we measured the area of several segments along the germband, we found that the incubation with SU5402 – but not with IWP-3 – reduced the size of all analyzed segments (*p* = 0.0034, *p* < 0.0001, *p* = 0.0007 and *p* = 0.0001 for Maxillary, Labial, Thoracic 1 and Thoracic 2 segments, respectively) (Fig 3D).

**Fig 3.**
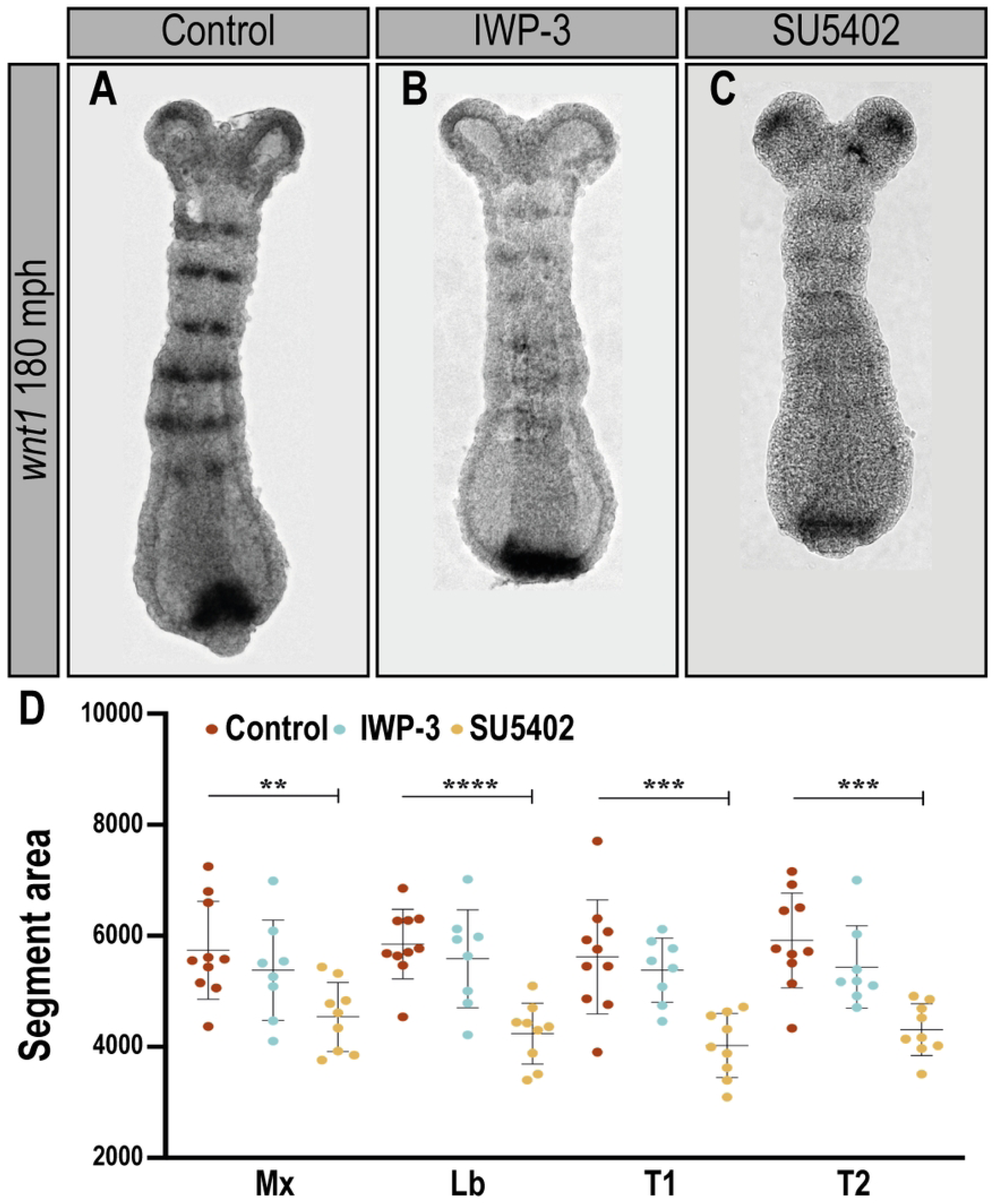
G**ermband segment size is reduced in Fgf signaling inhibited embryos.** (A-C) Representative pictures of 180 mph elongated germbands show their segments by *Tc-wnt1 in situ* hybridization. (A) DMSO (Control), (B) 45 µM IWP-3 and (C) 2.5 µM SU5402 treated embryos. (D) Segment area measurement after Control (*n* = 10), Wnt (*n* = 8), and Fgf (*n* = 9) inhibition treatments. Segments analyzed: Maxillary (Mx), Labial (Lb), Thoracic 1 (T1) and Thoracic 2 (T2). The error bars indicate the SD of the mean. Asterisks indicate statistically significant differences according to the Unpaired *t*-test. For SU5402 treatment, ** *p* = 0,0034 at Mx, **** *p* = < 0.0001 at Lb, *** *p* = 0.0007 at T1, and *** *p* = 0.0001 at T2.

**Fig 4.**
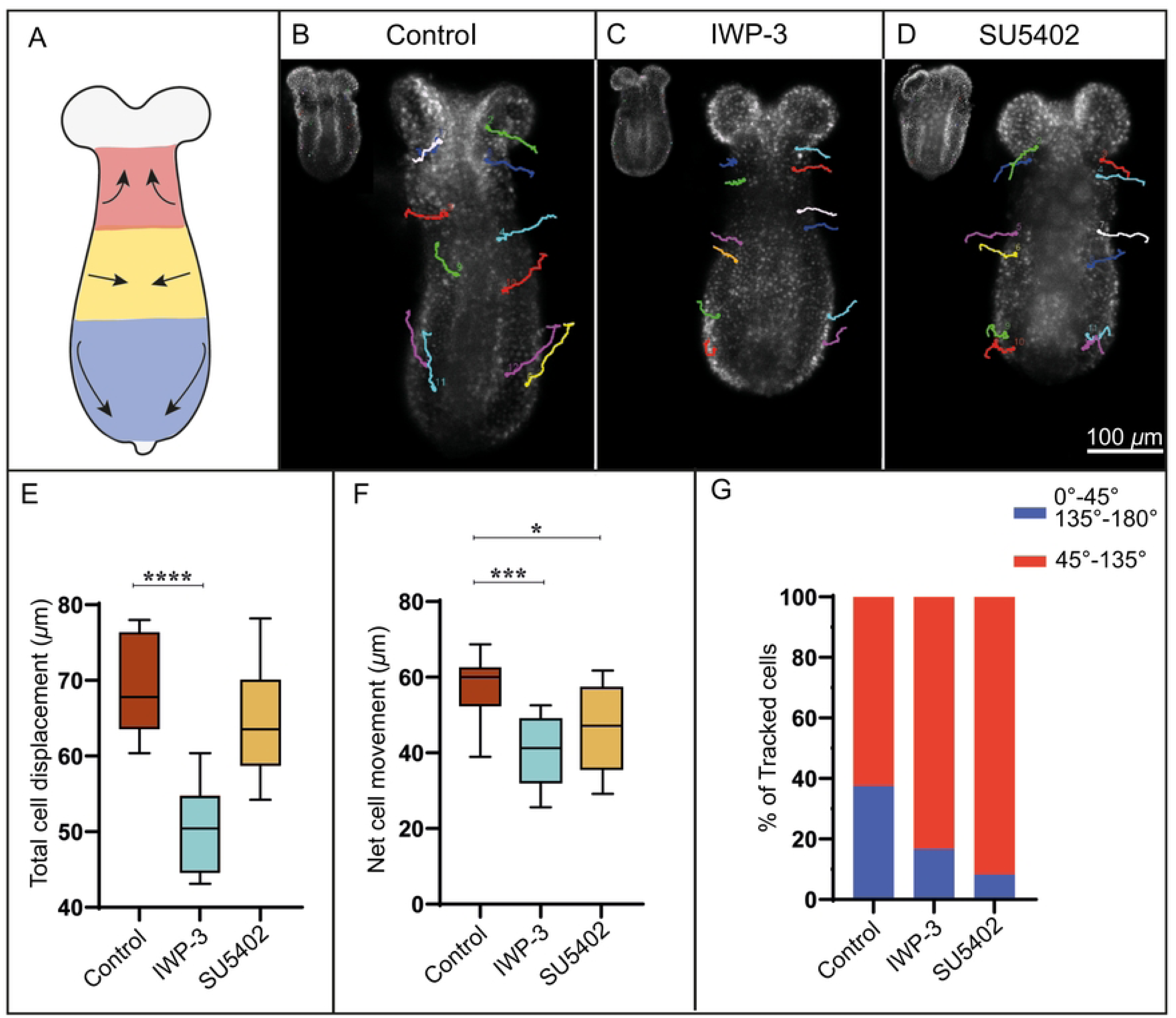
Wnt and Fgf signaling inhibition differentially alter cell movements during elongation. (A) A schematic diagram shows the three regions of the *Tribolium* germband that were tracked during cell movement analysis: Anterior, middle, and posterior (*n* = 12 cells per embryo). (B-D) Representative images of individual nuclei tracked (colored lines) in dissected EFA-nGFP transgenic line embryos (*n* = 3 embryos per treatment) over 2 hours at 30° Celsius, exposed to DMSO (B), IWP-3 (C), and SU5402 (D). A picture of each germband’s initial stage (t = 0) is shown at the left top of the image. Scale bar: 100 µm. (E-G) Various parameters obtained from the time-lapse analysis were evaluated, such as total cell displacement (E; *p* = 0.0001), net cell movement (F; *p* = 0.0004 for IWP-3 and *p* = 0.0262 for SU5402), and the orientation of cell movement (G). The resulting angles from cell movements were categorized as either in line (0 to 45° plus 135° to 180°) with or perpendicular (45° to 135°) to the elongation of the embryo (G). The error bars indicate the SD of the mean. Asterisks denote statistically significant differences as determined by a one-way ANOVA followed by a multiple comparison Tukey’s test.

### Wnt and Fgf signaling inhibition differentially alter cell movements during elongation

To address the potential involvement of the studied signaling pathways in the convergent and extension (CE) movements along the elongating *Tribolium* germband, we performed live imaging in dissected inhibitor- and DMSO/control-treated embryos (Fig 4). First, we mapped cell movements from three different regions of the embryo (anterior, middle, and posterior germband; see Fig 4A) over 2 hours by tracking individual fluorescent nuclei from an EFA-nGFP transgenic line (Fig 4B-D; *n* = 3 per treatment). Then, we could evaluate total cell displacement, net cell movement, and the resultant vector angle from the observed cell movements. Interestingly, these three parameters were differently affected in the presence of each inhibitor.

The total cell displacement, the sum of the distances covered by each cell during the analyzed period, was significantly reduced in the presence of the Wnt inhibitor. Specifically, the displacement was 69.23 µm in the control and 50.16 µm in the IWP-3-treated germbands (Fig 4E). This effect was observed throughout the embryo, with significant differences between the control and IWP-3 treatment present in the anterior, middle, and posterior regions (Anterior: Control = 63.21 µm, IWP-3 = 46.42 µm; Middle: Control = 69.84 µm, IWP-3 = 48.62; Posterior: Control = 74,64 µm, IWP-3 = 55.42 µm; S3 Fig). Interestingly, the inhibition of Fgf signaling did not significantly affect cell displacement but did have some impact on the net advances during elongation, as we will see below.

Furthermore, both inhibitors significantly reduced the net cell movement compared to controls, with IWP-3 showing a more significant impact than SU5402 when considering the whole embryo (Fig 4F). By region, there is a statistically higher effect on IWP-3-treated embryos at the anterior and posterior regions and on SU5402-treated embryos only at the posterior region compared to the control (S4 Fig).

Finally, the analysis of the resultant vector angle of the whole germbands showed a change in the orientation of the cell movements, where control embryos exhibit 37.5% of the tracked cells aligned with the anteroposterior axis, unlike IWP-3- and SU5402-treated germbands, that showed 16.7% and 8.3% of cells moving in an anteroposterior direction, respectively (Fig 4G). Analyzed by region, at the posterior end of the embryo, all SU5402-treated cells moved perpendicularly concerning the main axis, whereas 87.5% of control cells and 37.5% of IWP-3-treated cells moved along the AP axis (S5 Fig). The anterior region of control– and IWP-3-exposed embryos showed no cell movements aligned with the AP axis, while SU5402-treated embryos displayed 25% of cells moving in that direction (S5 Fig). Regarding the middle region, the percentages of cells moving along the anteroposterior axis were 25%, 12.5%, and 0% for control, IWP-3- and SU5402-treated germbands, respectively (S5 Fig).

## Discussion

Since *Tribolium castaneum* is amenable to gene knockdown by RNA interference (RNAi), most functional analyses have been accomplished in the offspring of female pupae or adult beetles injected with double-stranded RNA (pupal or parental/adult RNAi, respectively) [47, 48]. RNAi has become a robust genetic tool in *Tribolium* and many other organisms, allowing researchers to learn about the function of varied genes during development. *Tribolium* functional analysis by RNAi is mainly based on larval cuticle examination and embryonic gene expression patterns. On more than one occasion, only a few eggs are laid, producing embryos that display a range of phenotypes. Moreover, in some cases, we would like to avoid early effects obtained by using RNAi that could modify or disrupt a subsequent process of our interest. To circumvent these problems, we proposed using whole embryo culture for *in vivo* pharmacological manipulation [43]. Culturing dissected and/or bisected embryos allowed us to conduct functional analysis and compare gene expression under different conditions in the same embryo at different time intervals.

### Pharmacological inhibition of Wnt/Fgf signaling pathways

Given the prominent role that Wnt and Fgf signaling pathways have on the control of segmentation and body axis elongation in vertebrates [1, 49], several studies have evaluated their participation during arthropod elongation and segmentation [15, 16, 27, 30–33, 35, 36, 50–58]. To date, the establishment of the posterior zone, the control of the segmentation clock, and the segment border formation in arthropods are mainly attributed to the Wnt signaling [reviewed in 5, 18, 59]. Loss-of-function of different members of the Wnt pathway generates shorter embryos with defects in posterior segment formation. Most severe cases led to the complete elimination of segments after T2. In addition, many resulted in merged segments or the absence of segment boundaries. The first phenotype described is attributable to an early function of Wnt signaling, such as establishing the SAZ, and at least in *Tribolium*, possibly through regulating Fgf signaling [16]. Furthermore, segment boundary anomalies are probably related to the function of some ligands, such as Wnt1 in the beetle *Tribolium* and the spider *Parasteatoda* and Wnt8 in *Parasteatoda*, in setting up the segment polarity within each segment. Therefore, the pharmacological manipulation applied during germband extension allowed us to analyze the role of the Wnt pathway with the SAZ already formed and right during the elongation/segmentation process. Moreover, given the scarce experimental evidence suggesting a link between Wnt signaling and specific morphogenetic processes, such as cell division [35, 36, 57] and convergent extension [18], we make the most to evaluate these parameters during *Tribolium* elongation.

Our results showed that inhibiting Wnt and/or Fgf signaling pathways interfere with germband elongation, as was expected, mainly for Wnt. Interestingly, a significant reduction in cell divisions was found at the same concentrations that affected embryo lengthening for both drugs (45 µM for IWP-3 and from 2.5 µM to 50 µM for SU5402), suggesting a relationship between both processes, germband elongation and cell proliferation, as was shown before in Cepeda et al. [40], driven by these two signaling pathways. Given that germband lengthening and the spatiotemporal pattern of cell divisions are not uniform during *Tribolium* elongation [40, 42], we treated embryos with each inhibitor – and a combination of both – during different time intervals (1, 2, 3, and 4 hours after the onset of posterior elongation). We quantified the amount of PH3+ cells at each time point. Notably, all these reductions in the number of cell divisions were statistically more robust during a period characterized by its high rate of proliferation (120-180 mph), where a posteriorly localized peak of dividing cells shows up during elongation [40].

Even though the pattern of cell proliferation present in the beetle *Tribolium* is not shared between all studied arthropods [40], these results are in line with previous evidence in the spider *Parasteatoda tepidariorum* and the cockroach *Periplaneta americana,* which showed a decrease in cell proliferation after inhibiting Wnt signaling by RNAi and IWP-3 treatment [36, 57] or in the centipede *Strigamia maritima*, where its embryos displayed an increase in the size of the SAZ (possibly by a rise in cell divisions?) when exposed to LiCl, an activator of the Wnt pathway [53]. Added to this, the description made by Constantinou et al. (2020) [41] of a proliferating region within the SAZ of the branchiopod *Thamnocephalus platyurus* characterized by the expression of the Wnt4 ligand, agrees with the general conclusion that the Wnt pathway has a role, in addition to the maintenance of the SAZ and the establishment of the segment border, in the control of cell division during arthropod posterior growth.

The Fgf signaling pathway, much like the Wnt pathway, plays a critical role in vertebrate segmentation. It was the first signaling pathway to be experimentally linked to an undifferentiated posterior region, establishing a gradient of activity from posterior to anterior that defines the boundaries of newly formed segments during somitogenesis [60–62]. In experiments with chicken embryos, shorter segments were observed after *in ovo* electroporation of Fgf8 and after applying Fgf8-soaked beads [60]. Conversely, when Fgf signaling was inhibited through SU5402 treatment in zebrafish, longer somites were formed [62, 63]. Similar results were found in conditional Fgf receptor 1 mutant mouse embryos [64]. In contrast, our findings in Tribolium indicate that inhibiting Fgf signaling, unlike Wnt signaling, generates shorter segments.

Therefore, it appears that the effects of inhibiting Fgf signaling in the posterior growth of *Tribolium castaneum* are not directly linked to the segmentation process as seen in vertebrates. Instead, this may be related to the decrease in cell proliferation (evident from our results and those of [65]) or, more likely, to events associated with cellular mobility. In vertebrates, Fgf signaling has been linked to collective cell migration and cell motility in various biological systems [9, 66].

After inhibiting the FGF pathway, we noted a reduction in the net movement of cells compared to the control group, especially in the posterior region of the germband. Furthermore, in this region, the cell movements were predominantly oriented perpendicular instead of aligning with the main axis, significantly impacting the proper elongation of the embryo. These findings, along with a marked decrease in cell proliferation following exposure to SU5402, suggest that the FGF pathway, akin to the Wnt pathway, plays a crucial role in the axial elongation of *Tribolium* and likely other insects. This role is mediated by regulating cell division and cellular movement in the segment addition zone during posterior growth.

## Ackowledgments

We thank Rut Toro Díaz for her friendly technical assistance with the Cytation 5 Imaging Reader, from Universidad de Playa Ancha, Valparaíso. We also thank Constanza C. Macaya, Patricio E. Saavedra and Belén G. Riveros for their help at the very early stages of this work.

---

**s1 Fig.**
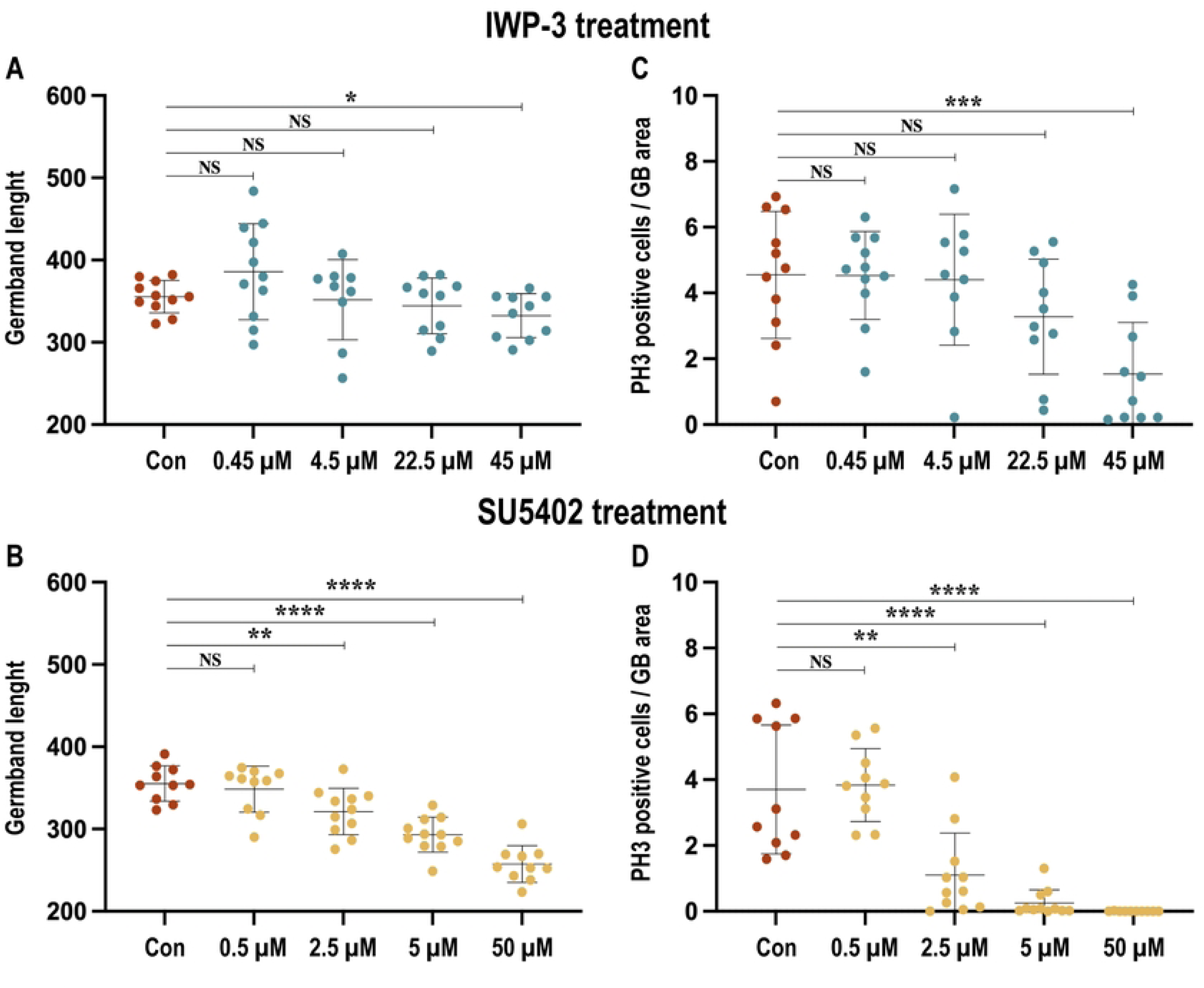

**s2 Fig.**
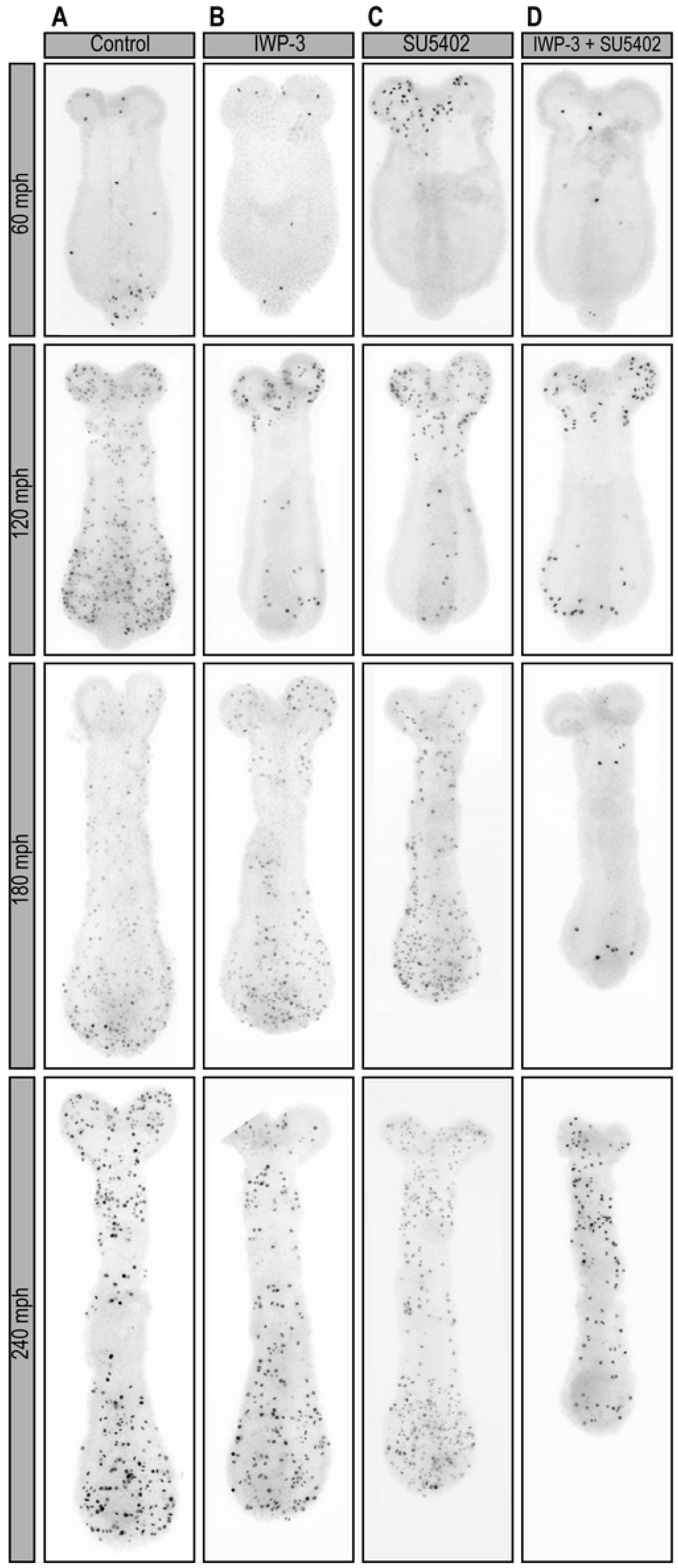

**s3 Fig.**
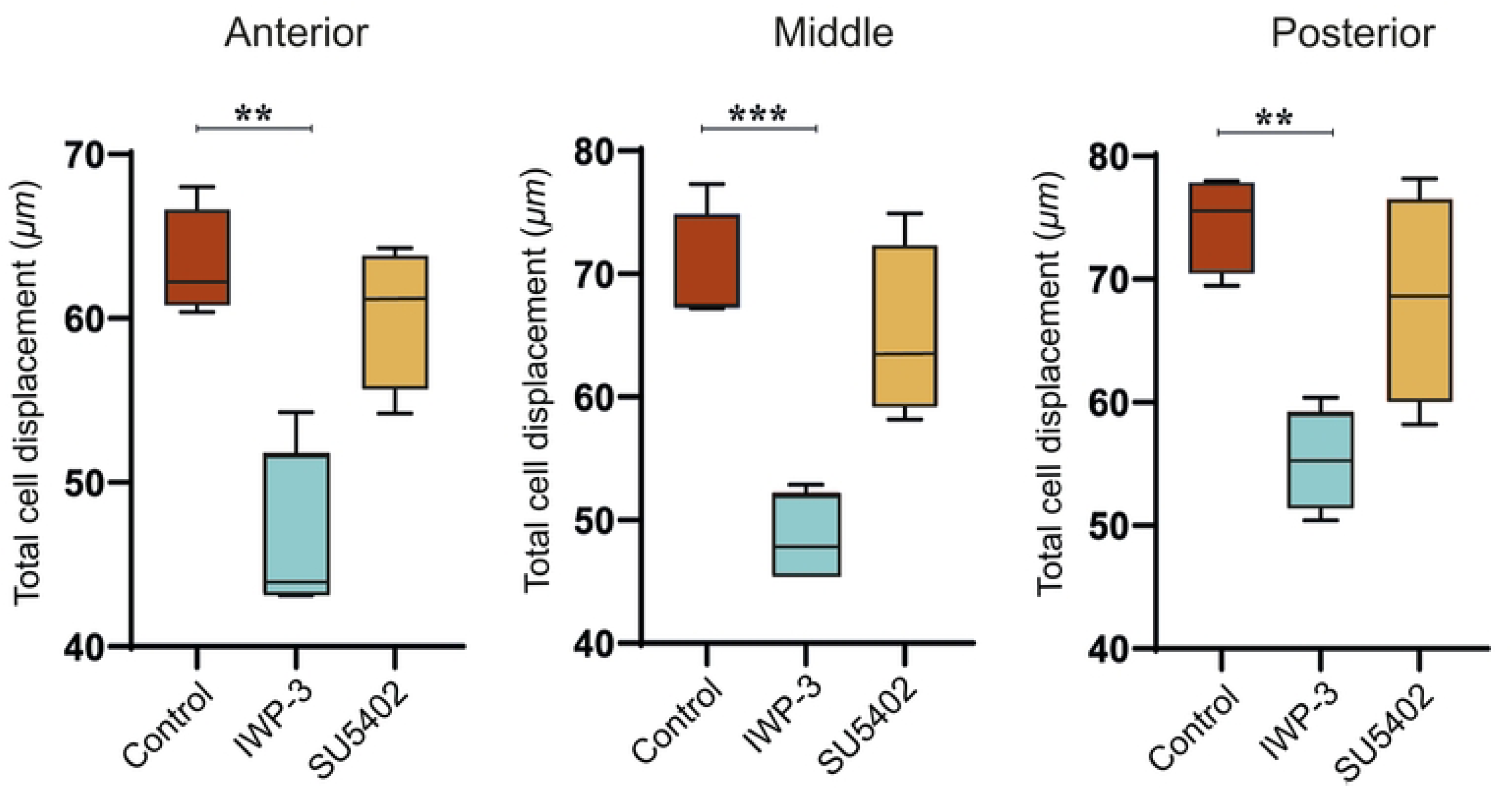

**s4 Fig.**
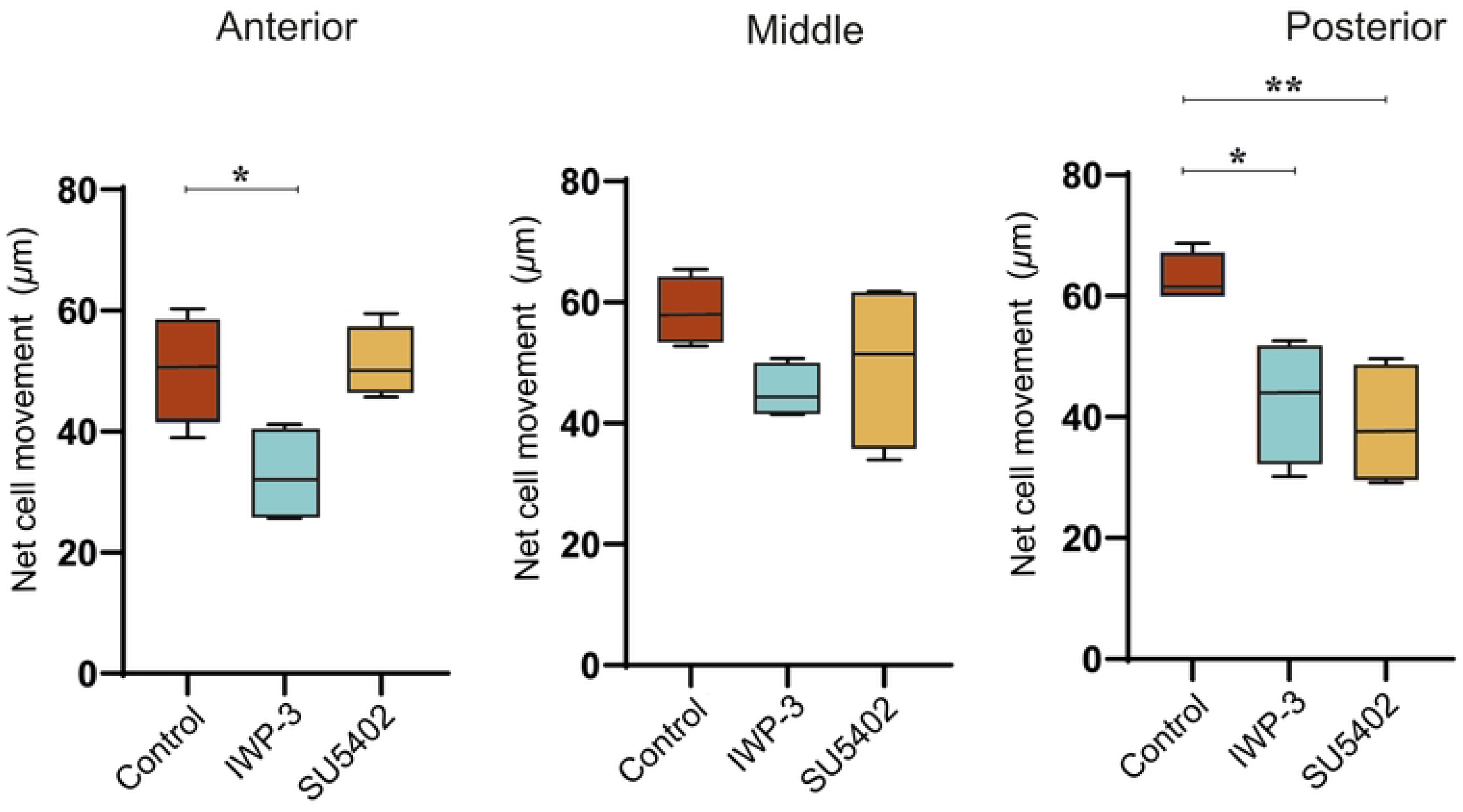

**s5 Fig.**
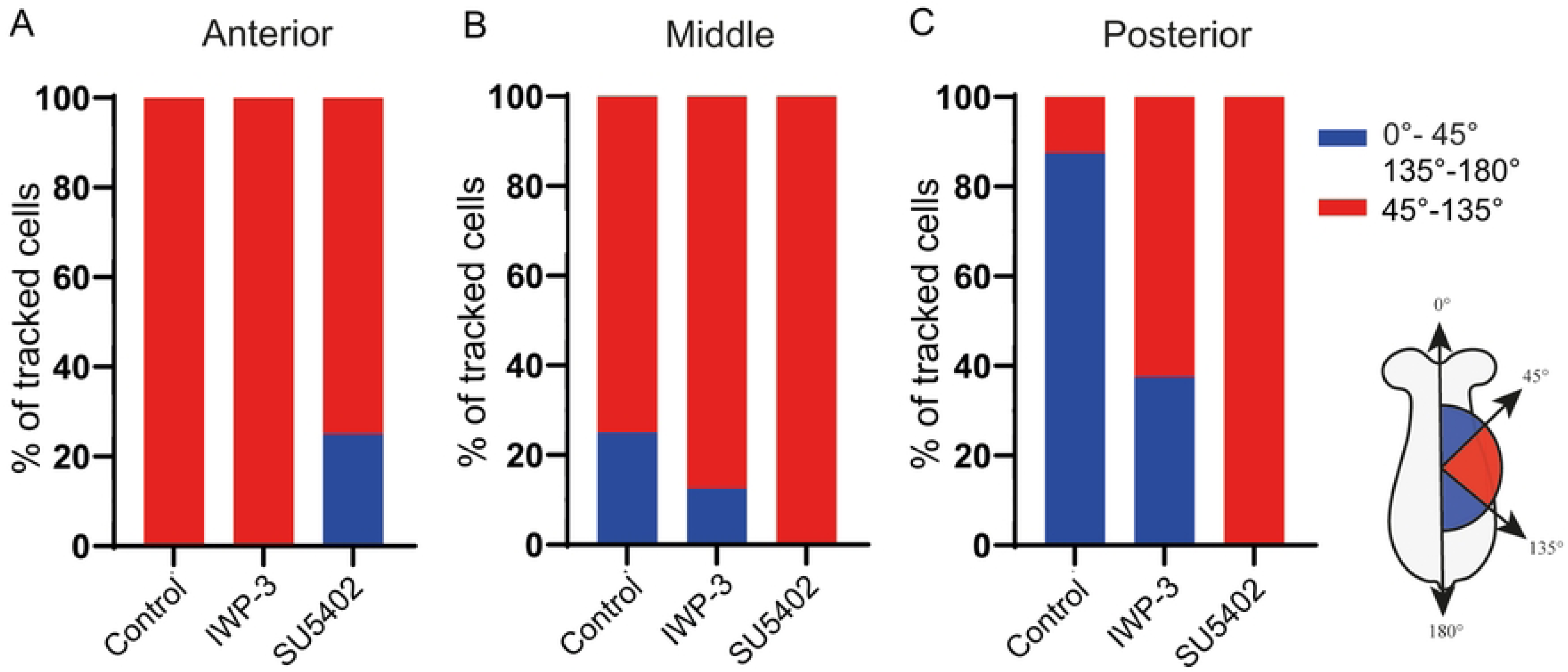

## Notes

### Competing Interest Statement

The authors have declared no competing interest.

